# A network-based approach to integrate nutrient environment in the prediction of synthetic lethality in cancer metabolism

**DOI:** 10.1101/2021.09.01.458495

**Authors:** Iñigo Apaolaza, Edurne San José-Eneriz, Luis V. Valcarcel, Xabier Agirre, Felipe Prósper, Francisco J. Planes

## Abstract

Synthetic Lethality (SL) is a promising concept in cancer research. A number of computational methods have been developed to predict SL in cancer metabolism, among which our network-based computational approach, based on genetic Minimal Cut Sets (gMCSs), can be found. A major challenge of these approaches to SL is to systematically consider tumor environment, which is particularly relevant in cancer metabolism. Here, we propose a novel definition of SL for cancer metabolism that integrates genetic interactions and nutrient availability in the environment. We extend our gMCSs approach to determine this new family of metabolic synthetic lethal interactions. A computational and experimental proof-of-concept is presented for predicting the lethality of dihydrofolate reductase inhibition in different environments. Finally, our novel approach is applied to identify extracellular nutrient dependences of tumor cells, elucidating cholesterol and myo-inositol depletion as potential vulnerabilities in different malignancies.

## Introduction

The main challenge of precision oncology is to be able to translate accumulating –omics data into actionable treatments, personalized for individual patients (Schüssler-Fiorenza Rose *et al*., 2019). Synthetic Lethality (SL) define as a type of genetic interaction where the co-occurrence of two (or more) genetic events results in cellular death, while the occurrence of either event on its own is compatible with cell viability represents a promising approach (Iglehart and Silver, 2009). Given the underlying genetic variations in tumor cells, SL largely expands the number of possible drug targets and creates an opportunity for more selective therapies (Rehman *et al*., 2010).

Extensive work has been done to predict SL in cancer using both experimental and computational approaches (Jerby-Arnon *et al*., 2014; Guo *et al*., 2016; McDonald *et al*., 2017; Carazo *et al*., 2019). These approaches have been mainly driven by the availability of large-scale gene knockout screening data for an increasing number of cancer cell lines (Tsherniak *et al*., 2017; Ghandi *et al*., 2019). Importantly, they provide an experimental *in vitro* measure of cancer gene essentiality, which can be integrated with genomic and transcriptomic data in order to hypothesize SL and identify response biomarkers.

Cancer metabolism is an ideal target to exploit the concept of synthetic lethality. Metabolic reprogramming of tumor cells leads to phenotypes that are substantially different from the ones observed in their healthy counterpart cells, and that can be potentially used to elucidate novel therapeutic strategies (Zecchini and Frezza, 2017). Outstanding works exploiting SL in cancer metabolism are reported in the literature (Frezza *et al*., 2011; Cardaci *et al*., 2015; Smestad *et al*., 2018), where different vulnerabilities were identified according to the underlying genetic context found in tumor cells.

We previously developed a computational framework to predict synthetic lethality in cancer metabolism based on the concept of genetic Minimal Cut Sets (gMCSs) (Apaolaza *et al*., 2017). Given a reference human genome-scale metabolic network, such as Recon3D (Brunk *et al*., 2018), gMCSs define combinations of gene knockout perturbations that block a particular metabolic target, in our case metabolites that are essential for cellular growth, *e*.*g*. nucleotides for DNA, amino acids for protein, lipids for cell membranes, etc. Using gene expression data as a proxy for the activity of metabolic enzymes, identified gMCSs were used to predict metabolic vulnerabilities in cancer. Our in-silico (network-based) approach was validated using large-scale *in vitro* gene-knockout screening data and *in vitro* functional studies in multiple myeloma.

Despite these promising results, the application of SL to cancer metabolism has still different challenges. One of them is the integration of tumor environment in the prediction of SL (Rancati *et al*., 2018). Certainly, the presence/deprivation of certain nutrients in the environment could modify the metabolic landscape and explain tumor resistance or sensitivity to metabolic targets (Halbrook *et al*., 2019; Tajan *et al*., 2021). However, this topic has not been systematically explored in current approaches to SL in cancer metabolism. In this article, we propose a new definition of SL that integrates tumor environment and extend our previous computational gMCSs approach to search for this family of synthetic lethal interactions. A computational and experimental proof-of-concept is presented for predicting the lethality in different environments of a well-studied drug target in cancer metabolism, dihydrofolate reductase (DHFR) (Vander Heiden, 2011). Finally, our novel approach is applied to predict extracellular nutrient dependences of *in vitro* cancer cell lines. We identified cholesterol and myo-inositol depletion as promising vulnerabilities of different tumors.

## Results

### Tumor nutrient environment and synthetic lethality

Figure 1 shows an example metabolic network under two different culture mediums (environmental contexts), CM1 and CM2, where metabolite C is essential for tumor growth. CM1 consists of a unique nutrient (M1) (Figure 1a). Under CM1, *g*_*1*_ and *g*_*2*_ form a synthetic lethal pair because their simultaneous inhibition blocks biomass production while individual inhibitions do not (Figure 1b). Note here that, if we consider a genetic context where *g*_*2*_ is not active (due to low expression, deletion or loss-of-function mutation, for example), *g*_*1*_ becomes an essential gene in this given context (Figure 1c).

**Figure 1.**
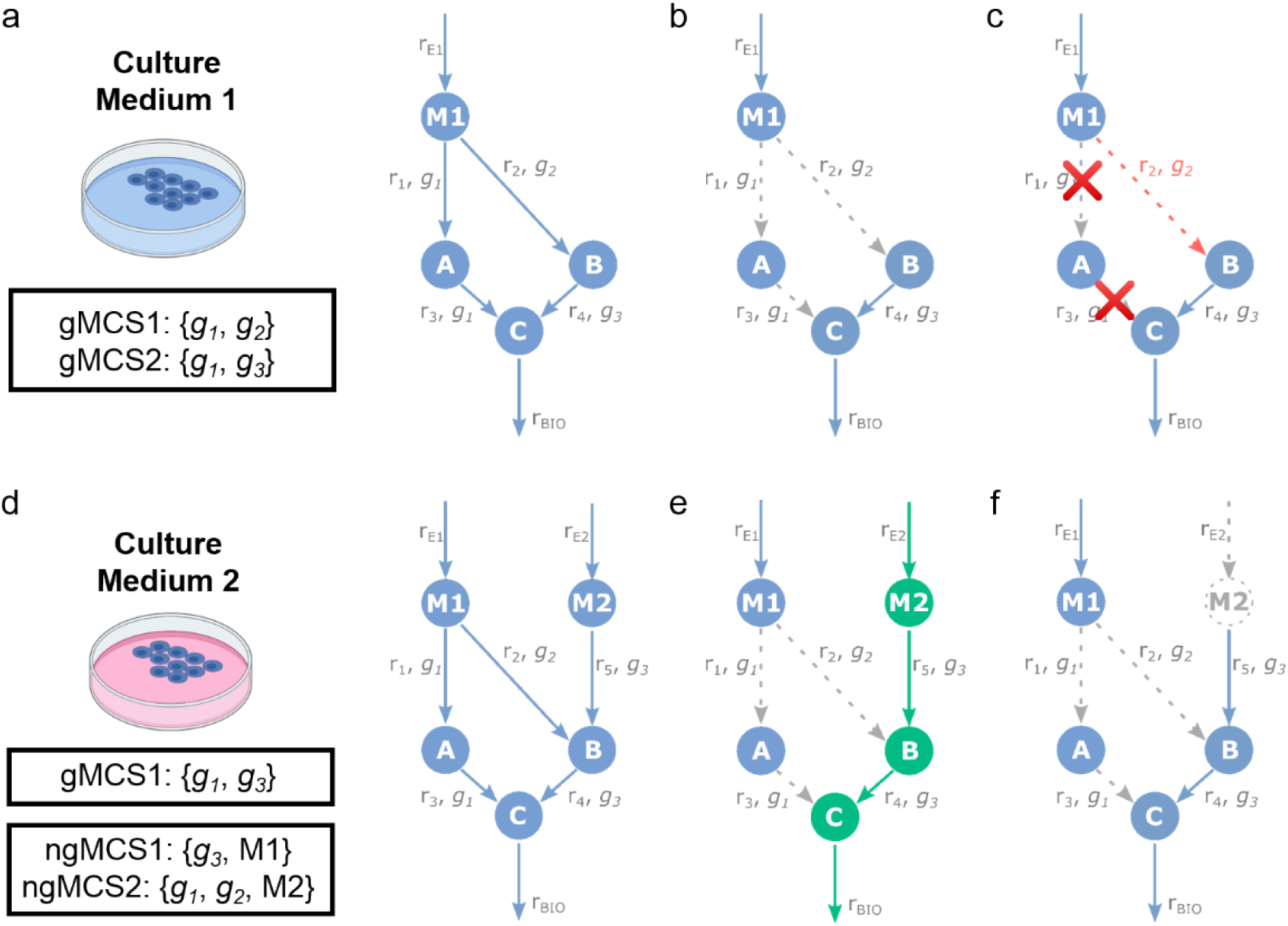
Synthetic Lethality in 2 different environmental contexts. **(a)** Example metabolic network under Culture Medium 1 (CM1). We have 5 reactions (r_E1,_ r_1_, r_2_, r_3_, r_4_), 4 metabolites (M1, A, B, C) and 3 genes (g_1_, g_2_, g_3_). r_E1_ is an input exchange reaction and represents the availability of M1. We have 2 gMCSs; **(b)** *g*_*1*_ and *g*_*2*_ are synthetic lethals under CM1. **(c)** If *g*_*2*_ is not active (red dotted line), the inhibition of *g*_*1*_ is essential under CM1. **(d)** Example metabolic network under Culture Medium 2 (CM2). With respect to the metabolic network in (a), we have 2 additional reactions (r_E2,_ r_5_) and 1 additional metabolite (M2). r_E2_ is an input exchange reaction and represents the availability of M2. We have 1 gMCS and 2 ngMCSs; **(e)** *g1* and *g2* are not synthetic lethal under CM2 due to the alternative pathway via M2 degradation (in green). **(f)** The inhibition of *g1* and *g2* and deprivation of M2 in the metabolic network from (d) renders cellular proliferation impossible. Gray dotted lines stand for not active/not present reactions/metabolites.

On the other hand, when trying to translate these findings to the network in CM2, due to the additional presence of M2 with respect to CM1 (Figure 1d), the simultaneous knock-out of the two genes (*g*_*1*_ and *g*_*2*_) is not lethal anymore (Figure 1e). This example illustrates that SL is dependent on the environmental context and a more general definition of SL is necessary in cancer metabolism in order to consider it together with the underlying genetic context. Here, we propose to identify synthetic lethal interactions involving genes but also nutrients in the environment.

In our example, Figure 1f shows an example of synthetic lethal involving two genes and one nutrient: {*g*_*1*_, *g*_*2*_, *M2*}, *i*.*e*. the lack of activity of *g*_*1*_ and *g*_*2*_ and the deprivation of nutrient *M2* leads to cellular death. In a particular context where *M2* is not present in the environment and *g*_*2*_ is not active, *g*_*1*_ remains essential. Similarly, in other context where *g*_*1*_ and *g*_*2*_ are not active, the depletion of *M2* is lethal, *i*.*e*. cellular growth is dependent on *M2* availability. Thus, this definition of SL is more general and allows us to identify both genetic and extracellular nutrient dependencies of tumor cells.

In order to be able to systematically identify this novel family of synthetic lethals, we extend our previous network-based approach, based on the genetic Minimal Cut Sets (gMCSs), leading to the concept of nutrient-genetic Minimal Cut Sets (ngMCSs). While gMCSs define minimal subsets of gene knockouts perturbations that lead to cellular death, ngMCSs incorporate nutrient deprivations as part of the predicted synthetic lethal interactions. Figure 1a,d show the list of gMCSs and ngMCSs for the 2 environments considered.

In genome-scale metabolic networks, nutrient deprivations can be modelled as the knockout of input exchange reactions, which represent the uptake of nutrients. In Figure 1, r_E1_ and r_E2_ are the input exchange reactions associated with the uptake of *M1* and *M2*, respectively. Thus, ngMCSs involve gene knockouts but also input exchange reaction knockouts. For example, ngMCS2:{*g*_*1*_, *g*_*2*_, *M*_*2*_} involves the gene knockouts of *g*_*1*_ and *g*_*2*_ and the reaction knockout of r_E2_, which is equivalent to the deprivation of *M2*. Our previously developed tool for the calculation of gMCSs, implemented in the COBRA toolbox (Heirendt *et al*., 2019), was amended to include the knockout of input exchange reaction in the solution space and calculate ngMCSs. Full details can be found in Methods section and Supplementary Data 1.

### Lethality of DHFR inhibition in different environments

As a proof-of-concept of our ngMCSs approach, we investigated a well-known metabolic target in cancer: dihydrofolate reductase (DHFR) (Luengo *et al*., 2017). The inhibition of DHFR has been proven lethal in different cancer cell lines under standard growth medium conditions (Zheng *et al*., 2018). Similarly, when we applied our gMCS approach to Recon3D (Brunk *et al*., 2018), a high-quality reference human genome-scale metabolic network, under RPMI growth medium conditions, the same result regarding DHFR essentiality was obtained (see Methods section). On the other hand, in a more complex tumor environment where all input nutrients annotated in Recon3D were available, we determined 17 gMCSs and 291 ngMCSs involving DHFR (Supplementary Data 2). We identified 2 ngMCSs that involve DHFR and exclusively nutrients not included in standard RPMI growth medium (ngMCS5 and ngMCS289 detailed in Supplementary Data 2). For illustration, one of them involves {DHFR, thymidine, dihydrothymine, thymine}. Since thymidine, dihydrothymine and thymine are not part of the RPMI growth medium, this ngMCS indicates the essentiality of DHFR under these *in vitro* conditions and support our ngMCS approach in a general context.

Among the list of ngMCSs mentioned above, we identified 2 nutrients which have been previously associated with resistance to DHFR inhibition, namely, thymidine and hypoxanthine (Zheng *et al*., 2018). In order to clarify our approach in a simpler environmental context, we re-calculated gMCSs and ngMCSs under the RPMI growth medium plus thymidine and hypoxanthine. Under this scenario, the list of gMCSs and ngMCSs involving DHFR is shown in Figure 2a. We have 2 ngMCSs, namely {DHFR, Thymidine} and {DHFR, Hypoxanthine}, which implies that the deprivation of either thymidine or hypoxanthine makes DHFR essential, as observed under standard RPMI growth medium. From a different angle, since DHFR is required in the de *novo* synthesis pathway of purines and pyrimidines, both thymidine and hypoxanthine are required to rescue proliferation upon DHFR knockout (Figure 2b). In addition, we have 4 gMCSs (Figure 2a), which indicate that the alternative salvage pathways through thymidine and hypoxanthine requires 1) the presence of the transporter SLC29A2 and 2) key enzymes, namely TK1 or TK2 for thymidine degradation, while HPRT1 or PNP or (GMPS and XDH) for hypoxanthine degradation (Figure 2b).

**Figure 2.**
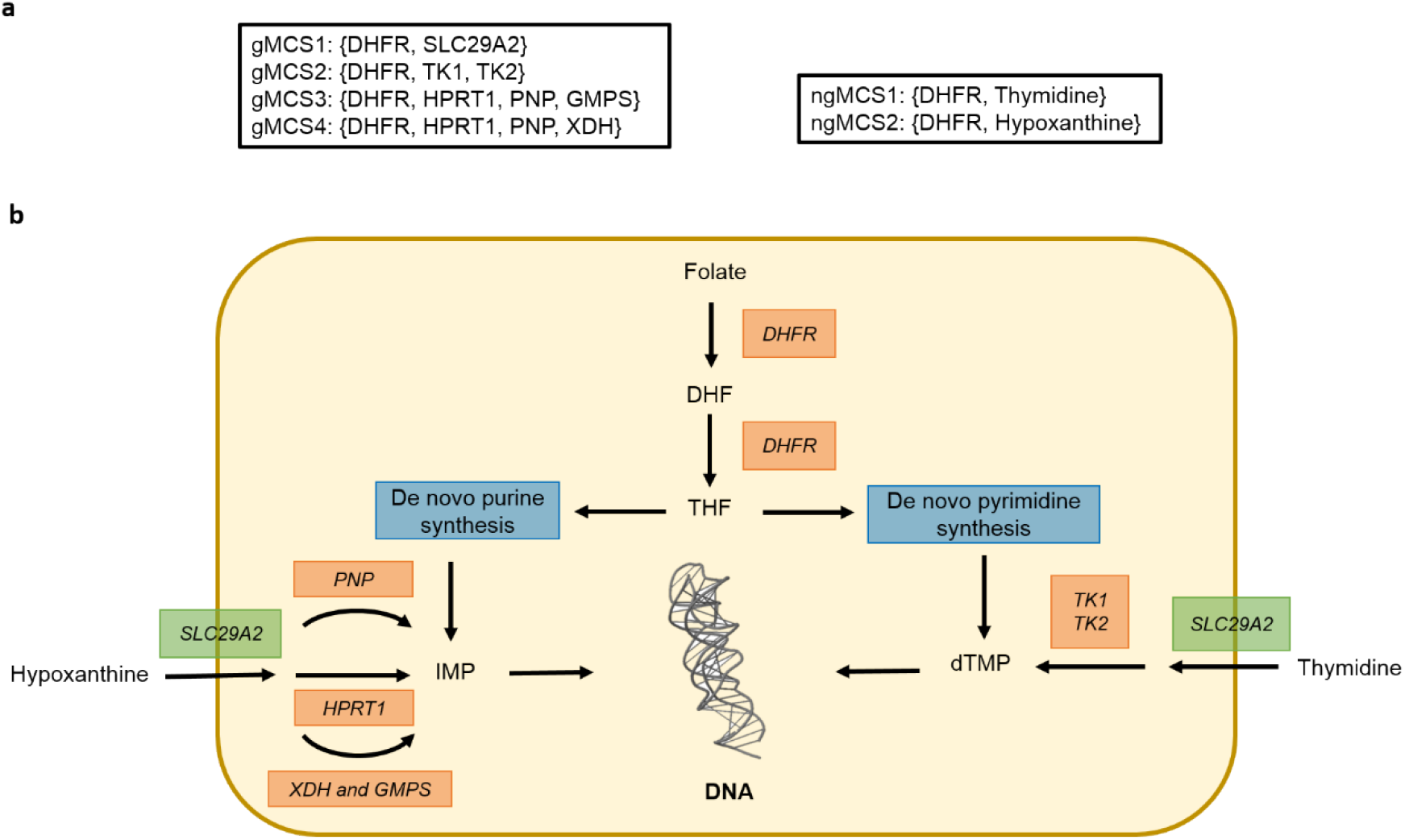
Predicted synthetic lethals involving DHFR with thymidine and hypoxanthine in the growth medium. **(a)** 4 genetic Minimal Cut Sets (gMCSs) and 2 nutrient-genetic Minimal Cut Sets (ngMCSs) involving DHFR. They were derived from Recon3D under the RPMI1640 growth medium plus thymidine and hypoxanthine; (**b)** Simplified network of metabolites and enzymes implied in the synthesis of purines and pyrimidines, emphasizing the role of DHFR, Thymidine and Hypoxanthine. Abbreviations: DHF: dihydrofolate; THF: tetrahydrofolate; IMP: inosinic acid; dTMP: 5-Thymidylic acid; DHFR: dihydrofolate reductase; PNP: purine nucleoside phosphorylase; HPRT1: hypoxanthine phosphoribosyltransferase 1; XDH: xanthine dehydrogenase; GMPS: Guanine Monophosphate Synthase; TK1: thymidine kinase 1; TK2: thymidine kinase 1; SLC29A2: solute carrier family 29 member 2.

To validate our hypothesis, we conducted experimental study in 3 different cancer cell lines: JVM2, HT29 and PF382. We used Methotrexate (MTX) for selective inhibition of DHFR. We first calculated GI50 for these cell lines and validated their sensitivity (IC50 < 50 nM) (Figure 3a). Second, we showed that the decrease in proliferation mediated by MTX is rescued when both thymidine and hypoxanthine are added into the growth medium (Figure 3b), in line with our computational predictions described above. A similar result was found when thymidine and hypoxanthine was added following *DHFR* silencing (Figure 3c), except for one of the siRNAs in JVM2 where no effect was observed. Note here that both *DHFR* siRNAs efficiently decreased DHFR expression in the three cell lines analyzed as detected by qRT-PCR (Fig. 3d). Finally, coherent gene expression of *SLC29A2, TK1* and *HPRT1* was observed in the three cell lines considered, according to CCLE (Cancer Cell Line Encyclopedia) (Ghandi *et al*., 2019) (see Supplementary Table 1), which enable alternative salvage pathways for purine and pyrimidine synthesis through thymidine and hypoxanthine, respectively.

**Figure 3.**
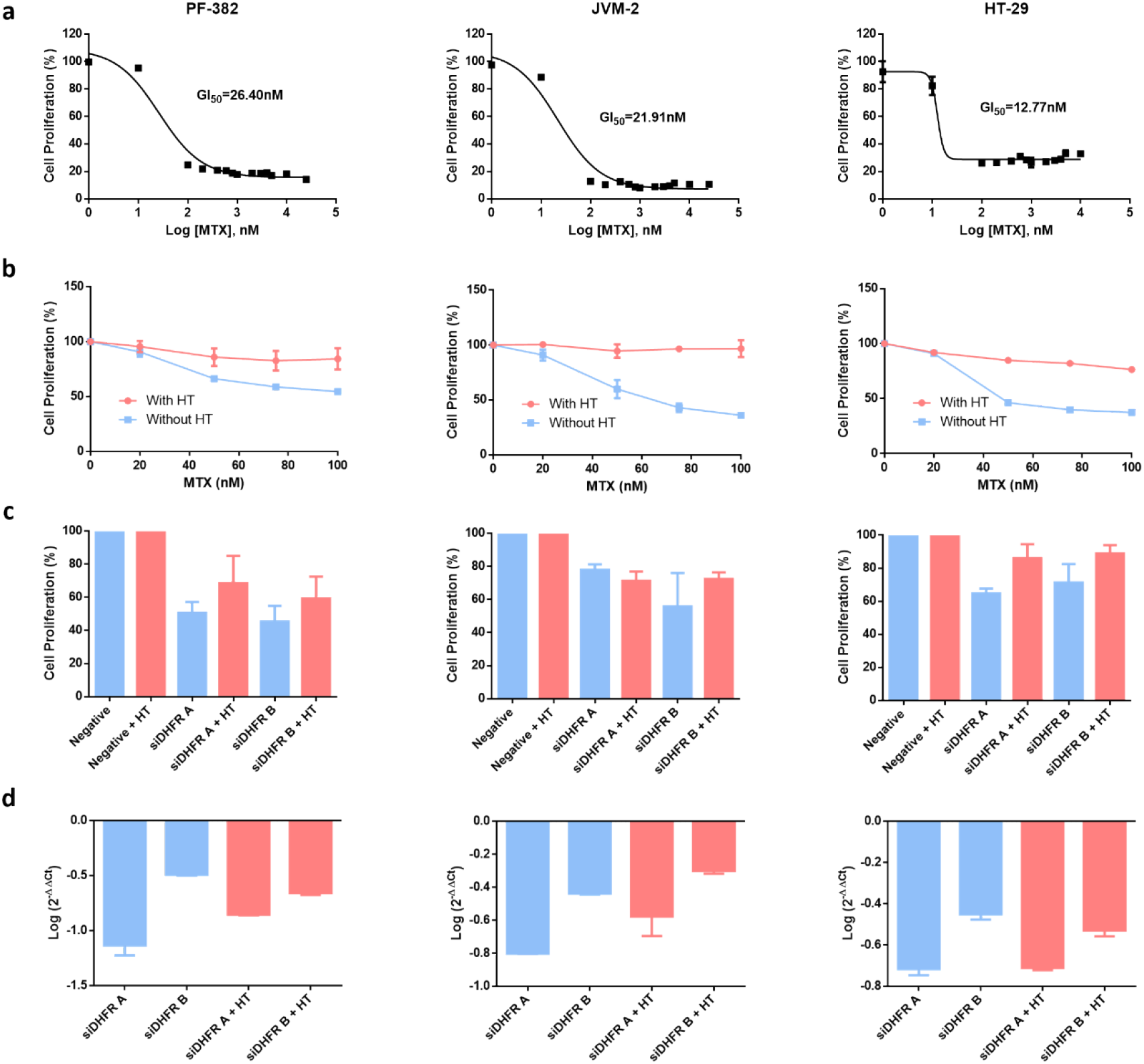
In-vitro experimental validation of DHFR inhibition with hypoxanthine and thymidine in the growth medium. **a)** GI50 values of MTX for PF-382, JVM-2 and HT-29 cell lines. **b)** Proliferation of PF-382, JVM-2 and HT-29 cell lines treated with different doses of MTX for 96h in presence or absence of Hypoxanthine-Thymidine. Data represent mean ± standard deviation of at least two experiments. **c)** Proliferation of PF-382, JVM-2 and HT-29 cell lines nucleofected with siRNAs targeted to *DHFR* gene in presence or absence of Hypoxanthine-Thymidine was studied by MTS at day 6 after nucleofection. The proliferation percentage refers to cells nucleofected with a negative control siRNA. Data represent mean ± standard deviation of at least two experiments. **d)** mRNA expression of DHFR gene 48 h after nucleofection with the specific siRNAs. Data are referred to GUS gene and an experimental group nucleofected with negative control siRNA. Data represent mean ± standard deviation of at least two experiments. MTX: methotrexate; HT: Hypoxanthine-Thymidine.

Considering the experimental data presented above, our ngMCSs approach was successful in determining in which environmental contexts DHFR inhibition is lethal in cancer. Based on our computational approach, the opposite case was explored below, *i*.*e*. which genetic context makes lethal an extracellular nutrient perturbation.

### Extracellular nutrient dependences of tumor cells

Our ngMCS approach can also be used to systematically identify context-specific nutrient dependences of tumor cells, *i*.*e*. supply of extracellular metabolites that are essential for tumor proliferation in a particular genetic context. To illustrate this, we searched for ngMCSs in Recon3D under RPMI growth medium conditions (see Methods section). We identified 41 ngMCSs (Supplementary Data 2), which involve 8 nutrients: L-asparagine, L-arginine, cholesterol, choline, D-glucose, L-glutamine, myo-Inositol and L-tyrosine. We discarded for further analysis choline and D-glucose because the associated genes in their ngMCSs show consistent and high expression and context-specific insights were not obtained. The opposite occurs with L-tyrosine, whose associated genes in the ngMCSs are lowly expressed in most cell lines. Thus, we focus on the other 6 nutrients (Figure 4a).

**Figure 4.**
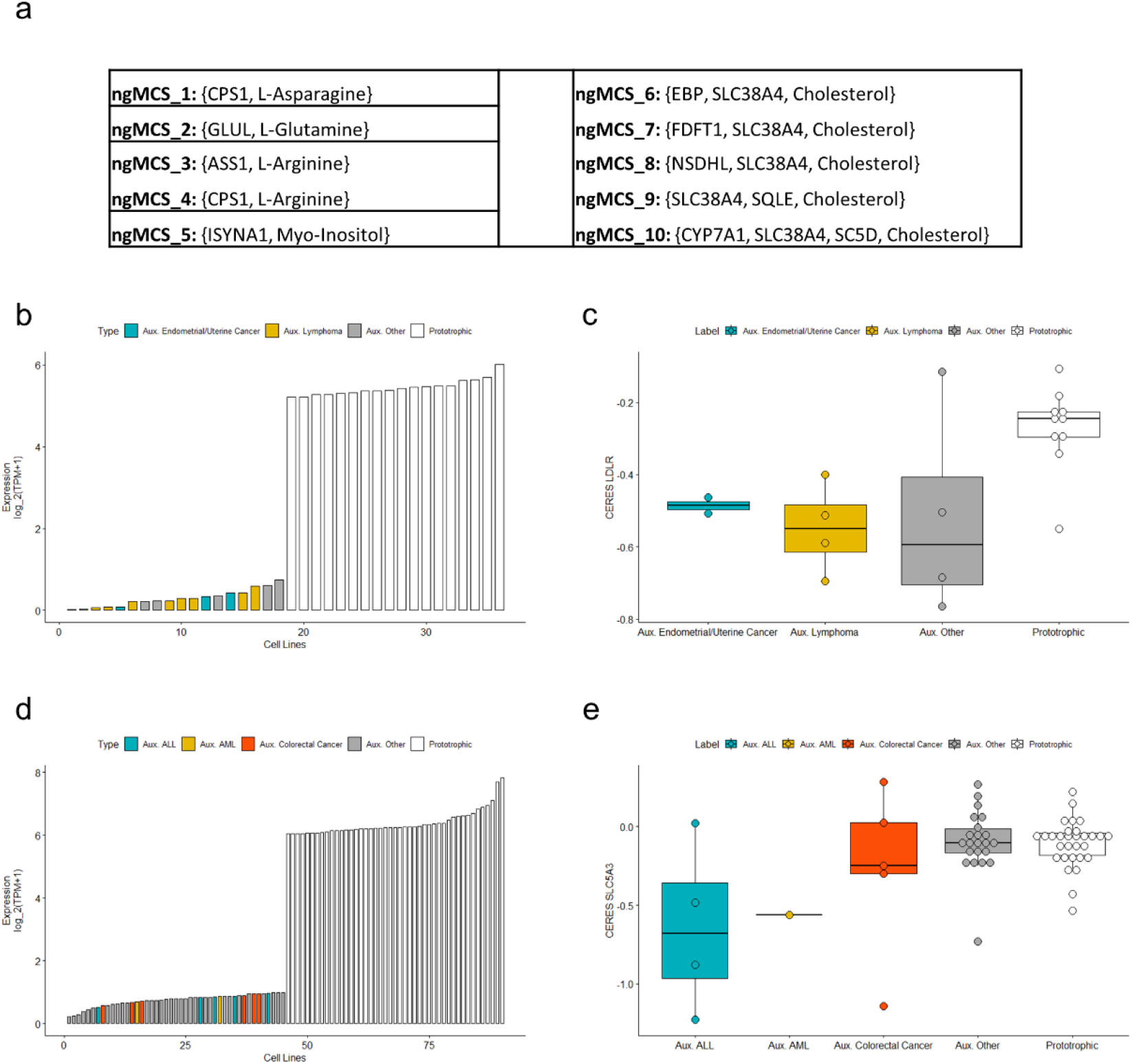
Extracellular nutrient dependences of *in vitro* cancer cell lines. **a)** Relevant ngMCSs involving L-Asparagine, L-Glutamine, L-Arginine, Myo-Inositol and Cholesterol. **b)** Expression levels of the most limiting gene in the ngMCSs associated with Cholesterol in 18 auxotrophic and 18 prototrophic cell lines. Auxotrophic cell lines were defined as those where the most limiting gene has an expression level below 1 TPM. For comparison, we took the same number of prototrophic cell lines; **c)** CERES scores for LDLR in the cell lines shown in b). The more negative CERES scores are, the more cellular proliferation will be decreased; **d)** Expression levels of the most limiting gene in the ngMCSs associated with Myo-Inositol in 45 auxotrophic and 45 prototrophic cell lines. **e)** CERES scores for SLC5A3 in the cell lines shown in d). Abbreviations: ‘Aux.’ refers to auxotrophic cell lines, *e*.*g*. Aux. ALL refers to auxotrophic ALL cell lines.

The dependency on extracellular L-asparagine in samples with loss of asparagine synthetase (ASNS) has been long studied in cancer research (Lazarus *et al*., 1969). In fact, L-asparagine depletion via asparaginase is an approved therapeutic strategy for acute lymphoblastic leukemia, and it is being investigated in solid tumors (Li *et al*., 2019). The dependence on extracellular L-arginine and L-glutamine in samples with loss of glutamine synthetase (GLUL) and Argininosuccinate Synthase 1 (ASS1), respectively, has been validated with *in vitro* experiments of nutrient depletion and barcode genetic screens (Li *et al*., 2019). The clinical importance of the uptake of L-arginine and L-glutamine in different tumors has received much attention in the last years (Zhang *et al*., 2017; Chalishazar *et al*., 2019).

In the case of L-arginine, we also identified the role of Carbamoyl-Phosphate Synthase 1 (CPS1), which is required for *de novo* biosynthesis of L-citrulline. L-citrulline is essential for *de novo* synthesis of L-arginine, but it is not present in the RPMI growth medium and, thus, its availability relies on *de novo* synthesis pathway via CPS1 and other genes (see Supplementary Data 2). CPS1 is a bottleneck in *de novo* synthesis of L-citrulline, being only expressed in intestinal epithelial cells and liver cells. Consequently, L-arginine is essential in most cases under RPMI growth medium unless L-citrulline is additionally supplemented, which was precisely the strategy followed in Li *et al*., 2019 in order to demonstrate the lethal interaction between L-arginine depletion and loss of ASS1. This insight again reinforces the importance of the tumor environment to predict synthetic lethality.

On the other hand, the dependence on extracellular cholesterol in tumors with loss of squalene monooxygenase (SQLE) was demonstrated in lymphoma (Garcia-Bermudez *et al*., 2019). They also showed that the inhibition of the low-density lipoprotein (LDL) receptor (LDLR) was a good proxy for cholesterol depletion. In addition to SQLE, our ngMCS approach in conjunction with RNA-seq data from CCLE (Ghandi *et al*., 2019), allow us to identify other genes implied in cholesterol auxotrophic cell lines: NSDHL, SC5D, FDFT1 and EBP. We identified 18 cell lines whose dependence on extracellular cholesterol was caused by the loss of one of these 5 genes, assuming a threshold of expression of 1 TPM. In agreement with the work of Garcia-Bermudez *et al*., 2019, 8 out of these 18 cholesterol auxotrophic cell lines are derived from lymphoma; however, we also detected 3 cell lines from endometrial adenocarcinoma (Figure 4b). For comparison, Figure 4b also includes 18 cell lines that are not dependent on extracellular cholesterol (cholesterol prototrophic cell lines), particularly those with the highest expression of the genes involved in the ngMCs associated with cholesterol. Using large-scale silencing data from DepMap (Tsherniak *et al*., 2017), we observed a similar effect of LDLR down-regulation in both lymphoma and endometrium cholesterol auxotrophic cancer cell lines, substantially superior than the one found in cholesterol prototrophic cell lines (Figure 4c).

In addition, patients with AML and loss of inositol-3-phosphate synthase 1 (ISYNA1) have shown a dependency on extracellular myo-Inositol (Wei *et al*., 2020). In this study, the inhibition of the myo-Inositol transporter SLC5A3 was presented as a good proxy for myo-Inositol depletion. Again, using RNA-seq data from CCLE, we identified 45 myo-Inositol auxotrophic cell lines, assuming a threshold of expression of 1 TPM for ISYNA1. Two of these cell lines are derived from AML; however, 5 and 6 of these cell lines correspond to acute lymphoblastic leukemia (ALL) and colorectal adenocarcinoma (CA) (Figure 4d). For comparison, Figure 4d also includes 45 cell lines that are not dependent on extracellular myo-Inositol, particularly those with the highest expression of ISYNA1. Using DepMap data, we observed that the most extreme effect of SLC5A3 down-regulation was found in ALL cell lines; however, we could not demonstrate a similar performance in colorectal cancer (Figure 4e).

## Discussion

There is an increasing body of literature evidencing that *in-vivo* resistance to metabolic vulnerabilities identified *in-vitro* could be mediated by alternative pathways driven by nutrients typically not included in standard growth media. This is illustrated here with our study about DHFR inhibition, whose anti-proliferative effect under RPMI growth medium is compensated with the addition of thymidine and hypoxanthine. There are other relevant cases in the literature. For example, it was recently reported that the supplementation of uridine rescues the anti-leukemic effect of dihydroorotate dehydrogenase inhibition (Sykes *et al*., 2016). A similar result was found in pancreatic ductal adenocarcinoma, where tumor-associated macrophages release pyrimidines to the extracellular medium which confer resistance to gemcitabine (Halbrook *et al*., 2019). On the other hand, restricting the availability of certain nutrients for tumor cells is an emergent strategy in cancer research. The dependence on L-asparagine in ALL is one paradigmatic example that is used clinically in patients with acute lymphoblastic leukemia. Other ongoing works include cholesterol depletion in lymphoma (Garcia-Bermudez *et al*., 2019) and myo-inositol depletion in AML (Wei *et al*., 2020), which illustrate that this therapeutic strategy could be exploited in wider settings. All together emphasizes the importance of systematically considering the nutrient environment to target cancer metabolism via synthetic lethality approaches (Muir and Vander Heiden, 2018).

Here, we propose a novel family of metabolic synthetic lethal interactions, which include genes but also nutrients in the environment, going beyond existing definitions in the literature. For their calculation, we extend our previously developed computational approach to identify synthetic lethal interactions in cancer metabolism (Apaolaza *et al*., 2017). In particular, we move from genetic Minimal Cut Sets, gMCSs, to nutrient-genetic Minimal Cut Sets, ngMCSs. While gMCSs define minimal subsets of gene knockouts perturbations that lead to cellular death, ngMCSs incorporate extracellular nutrient deprivation as part of the predicted synthetic lethal interactions. The ngMCS approach is a more flexible framework that allows us to predict context-specific genetic and nutritional perturbations that lead to cellular death.

Our computational tool could help to assess previously identified synthetic lethal interactions in more complex environmental scenarios and identify combinatorial therapies that additionally target alternative metabolic pathways. For example, based on the gMCSs shown in Figure 2, DHFR inhibition could be combined with SLC29A2 inhibition to avoid the uptake of hypoxanthine and disrupt the salvage pathway for purines biosynthesis. Our ngMCS approach could also predict nutrient restriction strategies that strengthen the efficacy of metabolic targets identified *in vitro*. This was illustrated in Figure 2, where restricting the availability of thymidine or hypoxanthine makes more effective the inhibition of DHFR in tumor cells. From another perspective, our approach opens new avenues to systematically identify novel response biomarkers to existing metabolic treatments, beyond frequently used genomic biomarkers (Setton *et al*., 2021).

In addition, based on RNA-seq data, the ngMCS approach can be used to predict extracellular nutrient dependences of tumor cells. In our analysis, summarized in Figure 4, we found that the lethality of cholesterol depletion previously reported in lymphoma can be extended to a subgroup of endometrial adenocarcinoma cell lines. Similarly, we pose the relevance of myo-Inositol depletion in acute lymphoblastic leukemia, going beyond previous results in acute myeloid leukemia. Although further work is required to assess the clinical relevance of these results, they illustrate the importance of studying more systematically the role of tumor environment in order to identify metabolic vulnerabilities.

The study of tumor environment and nutrient availability requires the use of metabolomic approaches. Availability of metabolomics data in different biofluids is growing day-by-day in cancer studies (Schraw *et al*., 2019); however, metabolomics studies in tumor extracellular microenvironment are less frequent but crucial to understand metabolic activity and identify metabolic vulnerabilities (Sullivan *et al*., 2019; García-Canaveras *et al*., 2019). Once this information becomes available for different tumors, our ngMCSs approach constitutes an elegant strategy to integrate genomics, transcriptomics and metabolomic data with genome-scale metabolic networks and predict synthetic lethals and metabolic vulnerabilities in cancer.

## Methods

### Metabolic Model

For the results presented above, we use Recon3D_3.01 as reference human genome-scale metabolic network (Brunk *et al*., 2018), which is available in https://www.vmh.life/. Recon3D_3.01 involves 13,543 reactions, where collectively participate 4,138 metabolites and 3,695 genes. Among them, we have a biomass reaction, which integrates essential metabolic requirements for cellular proliferation. The flux through the biomass reaction represents the proliferation rate, an important metabolic task in cancer studies. Note here that we corrected annotation errors previously identified in Recon 2 (Apaolaza *et al*., 2017; Pey *et al*., 2017) and inherited in Recon3D_3.01. In addition, we deleted HMR_9797 reaction, since adenine deaminase function has not been reported in human cells (Ribard *et al*., 2003). All corrections are summarized in Supplementary Table 2.

In the Results section, we used three different growth medium conditions. First, we simulated the RPMI1640 culture medium following the nutrient availability provided by the formulation of RPMI1640 with L-Glutamine (Lonza, Basel, Switzerland), similar to the one reported in Folger *et al*., 2011. Second, we simulated the most general growth medium conditions by enabling all input exchange fluxes of nutrients available in Recon3D_3.01. Finally, we simulated the RPMI1640 culture medium, as described above, plus thymidine and hypoxanthine, as discussed in Figure 2 and Figure 3.

Genome-scale metabolic networks can be used to predict metabolic synthetic lethals. In this approach, a synthetic lethal is defined as a subset of genes whoso simultaneous removal disrupts the flux through the biomass reaction. The identification of synthetic lethals was done through our previously developed approach termed genetic Minimal Cut Sets (gMCSs) (Apaolaza *et al*., 2017). gMCS define minimal subsets of gene knockouts perturbations that disrupt the flux through the biomass reaction. Here, we extend this concept in order to integrate the environmental context and nutrients availability. We described below how our previous formulation is amended to incorporate nutrient deprivation in the identification of synthetic lethal interactions, leading to nutrient-genetic Minimal Cut Sets (ngMCSs).

### Computation of nutrient-genetic Minimal Cut Sets

A central part of our gMCSs approach is the construction of the binary *G* matrix, where each row defines the reactions deleted by a minimal subset of gene knockouts. Then, based on duality theory and mixed-integer linear programming, we can identify minimal combinations of rows of *G* that blocks the biomass reaction. Full details can be found in Apaolaza et al. 2019 (Apaolaza *et al*., 2019).

In genome-scale metabolic reconstructions, input exchange reactions represent the availability of different nutrients in the environment. These reactions do not include any genetic association and, consequently, they cannot be blocked through any gene knockout, being their associated columns in *G* always zero. In order to consider their removal, which enables us to model the deprivation of nutrients in the environment, the *G* matrix must be amended. In particular, we need a new row for each nutrient in the environment with all entries equal to ‘0’ except for the column associated with its input exchange reaction. With this simple extension of *G*, ngMCSs, which are minimal strategies to block the biomass reaction through gene knockouts and/or nutrient deprivations, can be determined using our previously developed algorithms for gMCSs.

More specifically, in order to amend *G* matrix for the calculation of ngMCSs, we created an artificial gene for each different input exchange reaction present in the environment, *e*.*g*. gene_Ex_thymidine for the input exchange reaction of thymidine. This updated metabolic model was introduced as input data to our previously developed MATLAB function to calculate gMCSs (*CalculateGeneMCS*), freely available in the COBRA Toolbox (Heirendt *et al*., 2019), which requires IBM ILOG CPLEX to solve the underlying mixed-integer linear programming models. In this setting, the result could be both gMCSs and ngMCSs. We modified our MATLAB function to allow the user to calculate ngMCSs in the COBRA Toolbox (code available in Supplementary Data 1).

For the study of the lethality of DHFR, we followed two different strategies to calculate ngMCSs in the most complex environment with all nutrients present in Recon3D_3.01 available. First, we explored all possible combinations of gene knockouts and nutrient deprivations. Second, combinations of the DHFR knockout and nutrient deprivations were directly analyzed. Note here that *CalculateGeneMCS* function can restrict the search space among a predefined list of genes (*gene_set* optional parameter). The resulting gMCSs and ngMCSs are detailed in Supplementary Data 2. A similar analysis was done for the environment defined by the nutrients in the RPMI growth medium plus thymidine and hypoxanthine. The list of obtained gMCSs and ngMCSs are shown in Figure 2. For the study of extracellular nutrient dependencies of cancer cell lines, we considered all possible combinations of gene knockouts and deprivations of nutrients available in the RPMI growth medium. The list of ngMCSs can be also found in Supplementary Data 2. These results were computed with Intel(R) Xeon(R) Silver 4110 CPU @ 2.10GHz processors, limiting to 8 cores and 8 GB of RAM. A time limit of 60 seconds was set for each solution derived from the function *CalculateGeneMCS*.

### Computational modelling of DHFR knockout

The knockout of DHFR (dihydrofolate reductase) leads to the accumulation of DHF (dihydrofolate), which blocks the enzyme AICART (aminoimidazole carboxamide ribonucleotide transformylase) (Funk *et al*., 2013). At the same time, the inhibition of AICART increases the levels of 5-aminoimidazole-4-carboxamide-1-β-d-ribofuranosyl 5′-monophosphate (AICAR), which inhibits adenosine deaminase (ADA) and AMP deaminase (AMPDA) (Funk *et al*., 2013). In summary, in our presented analysis, the knockout of DHFR involves the removal of their associated reactions in Recon3D_3.01 and, indirectly, the reactions associated with AICART, ADA and AMPDA.

### Computational modelling of extracellular nutrient dependences in cancer cell lines

We neglected 5 reactions in Recon3D_3.01 annotated to protein degradation, since the metabolism of macromolecules is not well represented, leading to unbalanced and meaningless cycles. For consistency, we also deleted the availability of albumin from our simulations under the RPMI growth medium. This assumption does not affect the main results obtained for cholesterol and myo-Inositol.

### Cell culture

The cell lines PF-382 and JVM-2 were maintained in culture in RPMI1640 medium (Gibco, Grand Island, NY) and HT-29 cells with McCoy’s 5a medium (Gibco, Grand Island, NY), all of them supplemented with 10% fetal bovine serum (Gibco, Grand Island, NY) and penicillin/streptomycin (BioWhitaker, Walkersvill, MD) at 37 °C in a humid atmosphere containing 5% CO2. Cell lines were obtained from the DSMZ or the American Type Culture Collection (ATCC). All cell lines were authenticated by performing a short tandem repeat allele profile and were tested for mycoplasma (MycoAlert Sample Kit, Cambrex), obtaining no positive results.

### Cell proliferation assay

Cell proliferation was analyzed using the CellTiter 96 Aqueous One Solution Cell Proliferation Assay (Promega, Madison, W). This is a colorimetric method for determining the number of viable cells in proliferation. For the assay, cells were cultured by triplicate in 96-well plates, PF-382 at a density of 1×10^6^ cells/mL (100.000 cells/well, 100µL/well) and JVM-2 at a density of 2×10^5^ cells/mL (20.000 cells/well, 100µL/well). HT-29 cells were obtained from 80-90% confluent flasks and 100 µL of cells were seeded at a density of 5000 cells /well in 96-well plates by triplicate. Before addition of the compound, adherent cells were allowed to attach to the bottom of the wells for 12 hours. In all cases, only the 60 inner wells were used to avoid any border effects.

After 96 hours of treatment with different doses of methotrexate (MTX) (Selleckchem, TX, USA), plates with suspension cells were centrifuged at 800 g for 10 minutes and medium was removed. The plates with adherent cells were flicked to remove medium. Then, cells were incubated with 100 μL/well of medium and 20 μL/well of CellTiter 96 Aqueous One Solution reagent. After 1-3 hours of incubation at 37 ºC, the plates were incubated for 1-4 hours, depending on the cell line at 37 ºC in a humidified, 5 % CO2 atmosphere. The absorbance was recorded at 490 nm and 640 nm as a reference wavelength, using 96-well plate readers until absorbance of control cells without treatment was around 0.8. The background absorbance was measured in wells with only cell line medium and solution reagent. First, the average of the absorbance from the control wells was subtracted from all other absorbance values. Data were calculated as the percentage of total absorbance of treated cells/absorbance of non-treated cells. The GI_50_ values of the different compounds were determined using non-linear regression plots with the GraphPad Prism v5 software.

### MTX treatment

PF-382, JVM-2 and HT-29 cells were seeded by triplicate in 96-well plates at 100000, 20000 and 5000 cells per well. HT-29 cells were plated 24 hours before treatment. Then, all cell lines were treated with 20, 50, 75 and 100nM of MTX (Selleckchem, TX, USA) in presence or absence of 100μM hypoxanthine and 16μM thymidine (HT) (ThermoFisher Scientific). The cell proliferation assay was performed 96 hours after as described above. First, the average of the absorbance from the control wells was subtracted from all other absorbance values. Data were calculated as the percentage of total absorbance of treated cells/absorbance of non-treated cells.

### Cell transfection

Cells were passaged 24 h before nucleofection and culture was divided into two parts. One continu. One continues growing under same conditions (absence of HT) while the other was supplemented with 100μM hypoxanthine and 16μM thymidine. The transfection of siRNAs was done with the Nucleofector II device (Amaxa GmbH, Köln, Germany) following the Amaxa guidelines. Briefly, 1 × 10^6^ of PF-382 and JVM-2 cells were resuspended in 100 µL of supplemented culture medium, with our without HT, with 75 nM of *DHFR* siRNAs or Silencer Select Negative Control-1 siRNA (Ambion, Austin, TX) and nucleofected with the Amaxa nucleofector apparatus using programs C-009 and C-006, respectively. In the case of HT-29, siRNAs were transfected using lipofectamine transfection reagent 2000 (Invitrogen, Carlsbad, CA) according to manufacturer’s protocol. Briefly, HT-29 cells (50,000 cells per well) were seeded in a six-well plate with antibiotic-free medium 24 h before transfection. Cells were then incubated with transfection mixtures containing 75 nM of siRNAs or Silencer Select Negative Control-1 siRNA (Ambion, Austin, TX) for 4 h. Then, medium was replaced with full culture medium. Transfection efficiency was determined by flow cytometry using the BLOCK IT Fluorescent Oligo (Invitrogen Life Technologies, Paisley, UK). We used four different siRNAs against *DHFR* target, two for the isoforms 1, 3 and 4 and another two for the isoform 2 (siDHFR-134 A: AAGUCUAGAUGAUGCCUUA; siDHFR-134 B: AACCAGAAUUAGCAAAUAA; siDHFR-2 A: AGUACAAAUUUGAAGUAUA; siDHFR-2 B: AAAUUGAUUUGGAGAAAUA) to demonstrate that the results obtained with *DHFR* siRNA nucleofection are not due to a combination of inconsistent silencing and sequence specific off-target effects. Silencer Select Negative Control-1 siRNA was used to demonstrate that the nucleofection did not induce non-specific effects on gene expression. Nucleofection was performed twice with a 24 h interval. After 48 h of the second nucleofection, the *DHFR* mRNA expression was analyzed by qRT-PCR (*GUS* was employed as the reference gene). Cell proliferation was analyzed 0, 2, 4 and 6 days after two repetitive transfections as described above. HT were refreshed every two days. First, the average of the absorbance from the control wells was subtracted from all other absorbance values. Data were calculated as the percentage of total absorbance of *DHFR* transfected cells/absorbance of control cells.

### Quantitative RT-PCR

The expression of *DHFR* was analyzed by qRT-PCR in PF-382, JVM-2 and HT-29 cell lines. First, total mRNA was extracted with Trizol Reagent 5791 (Life Technologies, Carlsbad, CA, USA) following the manufacturer instructions. RNA concentration was quantified using NanoDrop Specthophotometer (NanoDrop Technologies, USA). cDNA was synthesized from 1 µg of total RNA using the PrimeScript RT reagent kit (Perfect Real Time) (cat. no. RR037A, TaKaRa) following the manufacturer’s instructions. The quality of cDNA was checked by a multiplex PCR that amplifies *PBGD, ABL, BCR*, and *β2-MG* genes. qRT-PCR was performed in a 7300 Real-Time PCR System (Applied Biosystems), using 20 ng of cDNA in 2 µL, 1 µL of each primer at 5 µM (DHFR F:5′-CCATTCCTGAGAAGAATCGAC-3′; DHFR R:5′-GGCATCATCTAGACTTCTGGAAA-3′; GUS F: 5′-GAAAATATGTGGTTGGAGAGCTCATT-3′; GUS R:5′-CCGAGTGAAGATCCCCTTTTTA-3′), 6 µL of SYBR Green PCR Master Mix 2X (cat. no. 4334973, Applied Biosystems) in 12 µL reaction volume. The following program conditions were applied for qRT-PCR running: 50 °C for 2 min, 95 °C for 60 s following by 45 cycles at 95 °C for 15 s and 60 °C for 60 s; melting program, one cycle at 95 °C for 15 s, 40 °C for 60 s and 95 °C for 15 s. The relative expression of each gene was quantified by the Log2(−ΔΔCt) method using the gene *GUS* as an endogenous control.

## Data availability

The authors confirm that the data supporting the findings of this study are available within the article and its supplementary material.

## Acknowledgements

This work was supported by the Minister of Economy and Competitiveness of Spain [PID2019-110344RB-I00], PIBA Programme of the Basque Government [PIBA_2020_01_0055], Elkartek programme of the Basque Government [KK-2020/00008], Instituto de Salud Carlos III (ISCIII) [PI16/02024, PI17/00701, PI19/01352, PI19/01352, PI20/01306], CIBERONC (Co-financed with European Union FEDER funds) [CB16/12/00489], ERANET program ERAPerMed [MEET-AML], MINECO Explora [RTHALMY], Cancer Research UK and AECC under the Accelerator Award Programme [C355/A26819] and Fundación Ramon Areces [PREMMAM]. L.V.V. was supported by PFIS [FI17/00297] award from Instituto de Salud Carlos III (ISCIII).

## Author contributions

X.A., F.P. and F.J.P. conceived this study. I.A, L.V.V. and F.J.P. developed the ngMCS approach while I.A and L.V.V. performed the computational analysis. E.S.J. and X.A. performed experiments. All authors wrote, read and approved the manuscript.

